# AlphaMate: a program for optimising selection, maintenance of diversity, and mate allocation in breeding programs

**DOI:** 10.1101/250837

**Authors:** Gregor Gorjanc, John M. Hickey

## Abstract

**Summary:** AlphaMate is a flexible program that optimises selection, maintenance of genetic diversity, and mate allocation in breeding programs. It can be used in animal and cross- and self-pollinating plant populations. These populations can be subject to selective breeding or conservation management. The problem is formulated as a multi-objective optimisation of a valid mating plan that is solved with an evolutionary algorithm. A valid mating plan is defined by a combination of mating constraints (the number of matings, the maximal number of parents, the minimal/equal/maximal number of contributions per parent, or allowance for selfing) that are gender specific or generic. The optimisation can maximize genetic gain, minimize group coancestry, minimize inbreeding of individual matings, or maximize genetic gain for a given increase in group coancestry or inbreeding. Users provide a list of candidate individuals with associated gender and selection criteria information (if applicable) and coancestry matrix. Selection criteria and coancestry matrix can be based on pedigree or genome-wide markers. Additional individual or mating specific information can be included to enrich optimisation objectives. An example of rapid recurrent genomic selection in wheat demonstrates how AlphaMate can double the efficiency of converting genetic diversity into genetic gain compared to truncation selection. Another example demonstrates the use of genome editing to expand the gain-diversity frontier.

**Availability:** Executable versions of AlphaMate for Windows, Mac, and Linux platforms are available at http://www.alpha-genes.roslin.ed.ac.uk/AlphaMate

## 1 INTRODUCTION

This paper describes the AlphaMate program that optimises selection, maintenance of genetic diversity, and mate allocation in breeding programs. Breeding programs aim to achieve defined targets over the course of a time horizon. Some programs select individuals to improve future performance, while other programs try to maintain the current state or even save a population from extinction. In all cases optimal management of genetic diversity within the bounds of practical constraints is crucial to sustainably support the current and yet unknown future targets. For example, breeding programs that select for improved performance must balance short and long-term genetic gain by avoiding excessive use of elite individuals. While elite individuals increase the mean of next generations, their excessive use also significantly reduces the amount of genetic diversity. This reduction limits the potential for long-term improvement. Breeding programs that focus solely on maintenance of diversity must also ensure that individuals contribute in a somewhat balanced manner. Therefore, breeding programs must balance individuals’ contributions to future generations to ensure long-term viability.

The optimal contribution theory formulates balancing selection and maintenance of genetic diversity as optimisation of individuals’ contributions to the next generation under constrained rate of group coancestry; see Woolliams et al. (2015) for review. Contributions can be optimised with two approaches. The first approach optimises contributions to maximise genetic gain under a constrained rate of group coancestry amongst the contributors or to only minimise group coancestry. This optimisation prevents the loss of genetic diversity above the accepted rate of coancestry. Optimisation of contributions can be followed by mate allocation to minimize inbreeding of individual matings. This second optimisation prevents excessive inbreeding depression in resulting progeny. These two optimisations can be solved with deterministic optimisation methods that vary according to the mathematical formulation of the problem, e.g., Lagrangian multipliers (Meuwissen, 1997), linear programming (Toro and Perez-Enciso, 1990), or quadratic programming (Pong-Wong and Woolliams, 2007). The second approach jointly optimises contributions and mate allocations via optimisation of a mating plan (Kinghorn and Shepherd, 1999; Akdemir and Sanchez, 2016). The joint optimisation does not have an analytical form and has to be solved with stochastic or metaheuristic methods, such as evolutionary algorithms. These methods can easily accommodate constraints and multiple objectives in comparison to deterministic algorithms, but usually require more computing time.

Existing programs that implement the above described approaches are often applicable to specific applications and are not generically applicable to both animal and plant populations or do not accommodate application of modern biotechnologies such as genome editing. The aim of this work is to present a flexible program AlphaMate that can be used in animal and cross- or self-pollinating plant populations. We briefly describe the implemented methodology in AlphaMate and show its application in two examples: i) maximising efficiency of converting genetic diversity into genetic gain in a rapid recurrent genomic selection program for wheat and ii) expanding the gain-diversity frontier with genome editing.

## 2 METHOD

AlphaMate by default jointly optimises contributions and mate allocations. The goal of this optimisation is to find a valid mating plan that delivers desired targets. This is achieved with an evolutionary optimisation of a single objective or multiple objectives simultaneously.

A valid mating plan is defined by a combination of mating constraints: i) the number of matings, ii) the maximal number of parents, iii) the minimal, equal, or maximal number of contributions per parent, or iv) allowance for selfing. If a user requires only optimised contributions, then only the specifications ii) and iii) are relevant.

The desired targets formulate optimisation objectives, such as: i) maximize genetic gain, ii) minimize group coancestry amongst contributors, iii) minimize expected inbreeding of individual matings, iv) maximize genetic gain with constrained group coancestry or inbreeding, or v) as i) or iv) but with the ability to genome edit a fixed set of contributors.

Optimisation is performed with an evolutionary algorithm based on differential evolution (Storn and Price, 1997) with modifications to avoid premature convergence (Gondro and Kinghorn, 2009). For a single target, we optimise a single objective function accounting for mating constraints. For multiple targets, we perform multiple objective optimisation in two steps; see e.g. Deb (2014) for review. First, we optimise single objective functions for each target separately to find bounds of the objective space and normalize objectives. Second, we use the ε–constraint method to either: i) find a Pa-reto-optimal solution with targeted balance between objectives or ii) evaluate the whole frontier of Pareto-optimal solutions (the Pareto frontier). A Pareto-optimal solution is the best solution with a specific balance between objectives. The Pareto frontier is a set of Pa-reto-optimal solutions and is useful when a breeder does not have clearly defined targets and can explore optimal solutions with different balance between targets to reach a decision. Fig. 1. demonstrates the Pareto frontier of genetic gain and group coancestry and the optimisation path for a targeted solution.

**Fig. 1.**
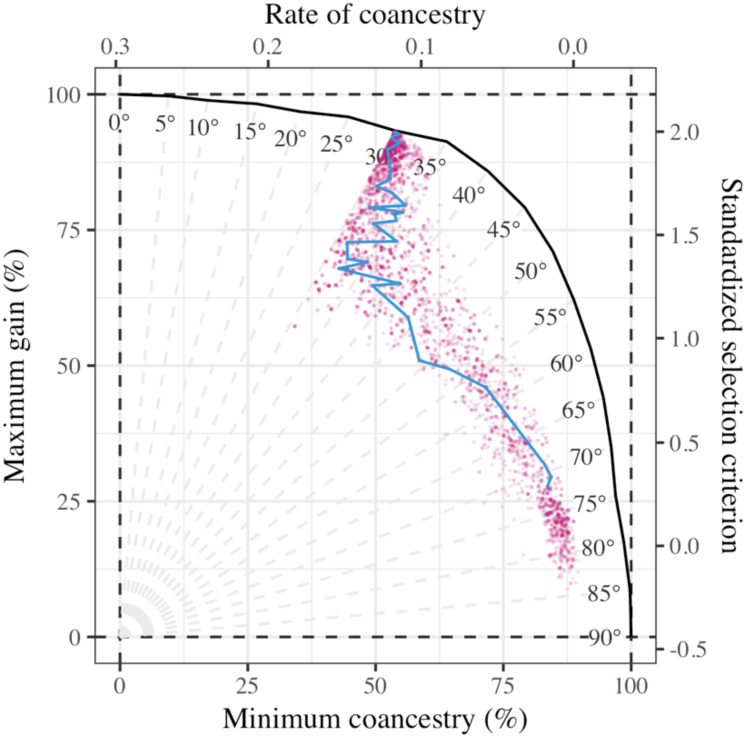
Trade-off between genetic gain and group coancestry and optimisation path of evolutionary algorithm (target set to 30°, dots show evaluated solutions, line shows evolution of the best solution)

Optimisation works with mating plans, which we encode as proposed by Kinghorn and Shepherd (1999). We ensure that mating plans are valid in two ways. First, we fix encoded representation, e.g., we trim contributions to user defined limits and round them to integer values (Lampinen and Zelinka, 1999). Second, when fixing is not sufficient, we penalize invalid mating plans so that the evolutionary algorithm advances (more) valid mating plans.

## 3 USE

AlphaMate was written in object oriented Fortran 95 as a standalone program and compiled versions are available for Windows, Mac, and Linux platforms. A single specification file controls the program. In this file, a user specifies: i) input files, ii) mating constraints, iii) desired targets, and iv) optimisation controls. Below we briefly describe these groups of specifications, while the full list is available in the AlphaMate manual.

i) The basic files are the coancestry matrix, selection criteria, and gender information for candidates. The coancestry matrix and selection criteria can be based on pedigree or genomic data. Additionally, further individual or mating specific information can be given to enrich optimisation objectives.

ii) Mating constraints can be gender specific or generic to accommodate different reproductive systems in animals and cross- or self-pollinating plants. A user can specify all the mating constraints or a subset of them depending on the objective of optimisation and biologic or logistic reasons.

iii) Desired targets define the optimisation objectives. For ease of use we allow for various forms of some targets, e.g., constraint on the loss of genetic diversity can be defined with the targeted value of coancestry, rate of coancestry, percentage of the minimum possible coancestry, or trigonometric degrees between genetic gain and group coancestry (see Fig. 1.).

iv) Optimisation controls specify weights used to combine multiple targets into a single objective function, penalties used to penalize invalid mating plans, and parameters of evolutionary algorithm such as the number of iterations, the number of evaluated matings plans, convergence criteria, etc.

The AlphaMate output consists of: i) summary of input data, ii) list of contributors with associated data and optimised contributions, iii) optimised mating plan, iv) optimisation log, and v) the seed value for random number generator to enable reproducibility. A utility R script is provided to plot the Pareto frontier and the optimisation paths.

## 4 DEMONSTRATION

We demonstrate the use of AlphaMate with two examples. The first example optimises conversion of genetic diversity into genetic gain based on a subset of the results from a previous study we undertook to model the benefit of rapid recurrent genomic selection in wheat (Gorjanc et al., 2017). Here we compare AlphaMate to truncation selection method over 20 years when breeding program used four recurrent selection cycles per year. In each cycle, we used a pool of 32 parents to generate 16 crosses with 160 progeny in total. We used AlphaMate to optimise selection and mate allocation with a constraint that a parent could only contribute at most four crosses. We supplied AlphaMate with genomic estimates of breeding values and a genomic coancestry matrix that measured as the proportion of marker alleles in common between the progeny. We ran ten simulations, collected genetic mean and genic standard deviation in progeny for each year, and fitted linear regression on this data. In Fig. 2. we show the evolution of genetic mean and genic standard deviation over the 20 years as influenced by different balances between selection and maintenance of genetic diversity achieved via different trigonometric degrees. We also show results for the truncation selection method, where we ignored maintenance of genetic diversity and parents either contributed one or four crosses. There is a clear effect of balancing the two objectives on the long-term performance of the breeding program. In comparison to truncation selection with one (four) cross per parent AlphaMate with the target of 35° delivered 65% (11%) higher genetic gain with 278% (139%) lower reduction of genic standard deviation, which translates to a 242% (93%) higher efficiency of converting genetic diversity into genetic gain. We note that truncation selection with one cross per parent achieved slightly higher genetic gain than AlphaMate with comparable efficiency (15-20°), which suggests that group coancestry based on the proportion of shared marker alleles might not be the best metric for the long-term maintenance of genetic diversity in populations under selection. This is the subject of our future research.

**Fig. 2.**
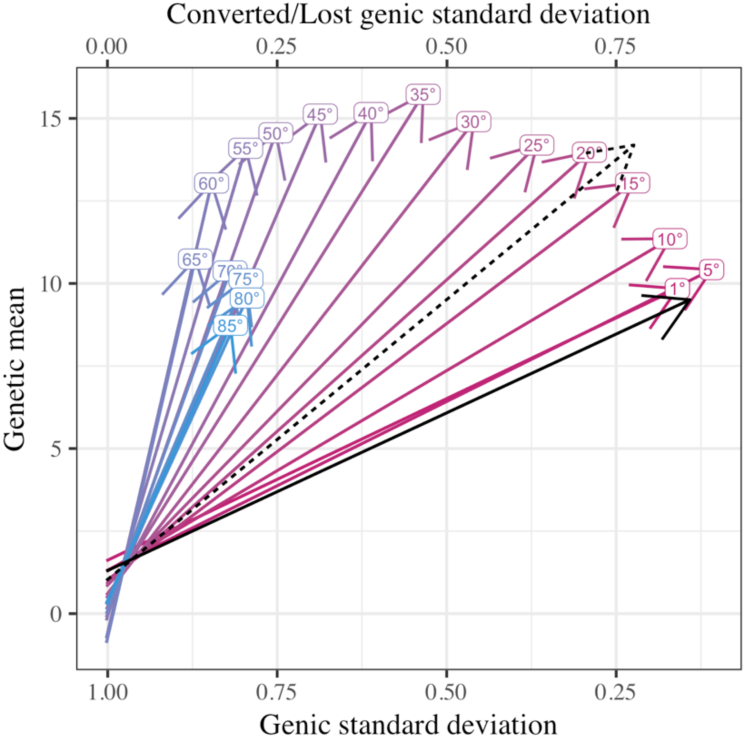
The genetic mean and genic standard deviation over 20 years of a wheat breeding program optimised with AlphaMate for different balance between selection and maintenance of genetic diversity defined by trigonometric degrees; black lines denote truncation selection with one (dashed) or four (full) crosses per parent

The second example expands the gain-diversity frontier based on our previous modelling of the genome editing potential to improve quantitative traits along standard selection methods (Jenko et al., 2015). By way of example genome editing could improve the genetic merit of the top individuals or the average individuals. If used optimally, the latter option might have the potential to expand the gain-diversity frontier, i.e., expand the Pareto frontier of genetic gain and group coancestry. To test this, we have simulated a breeding program as in Jenko et al. (2015) with 1000 selection candidates out of which we aimed to select 25 males and all 500 females with equalized contributions. In addition, we assumed to have resources to genome edit any 5 males, each at 1, 5, or 20 top causal loci. The question in such a setting is, which males should be selected and edited to maximise genetic gain for a given increase in group coancestry. We evaluated this by first calculating the genetic merit that male candidates could have been achieved with editing. We then provided the non-edited and edited genetic merit of the candidates to AlphaMate and jointly optimised which males should be selected and edited. To this end we have added to optimisation a set of “edit rank” variables of length equal to the number of candidates for editing. When calculating the genetic gain, we used “edited” genetic merit for individuals with the highest “edit rank” and “non-edited” genetic merit for the others. In Fig. 3. we show the Pareto frontier without and with genome editing. The results show that genome editing expanded the frontier. However, the expansion was substantially only when we edited 20 top causal loci and when target was not solely on minimum coancestry. At 30° the baseline maximum gain was 80% and the baseline minimum coancestry was 46%. With editing 5 or 20 loci the maximum gain improved to respectively 85% or 96%, while the minimum coancestry only slightly deteriorated to 45%.

**Fig. 3.**
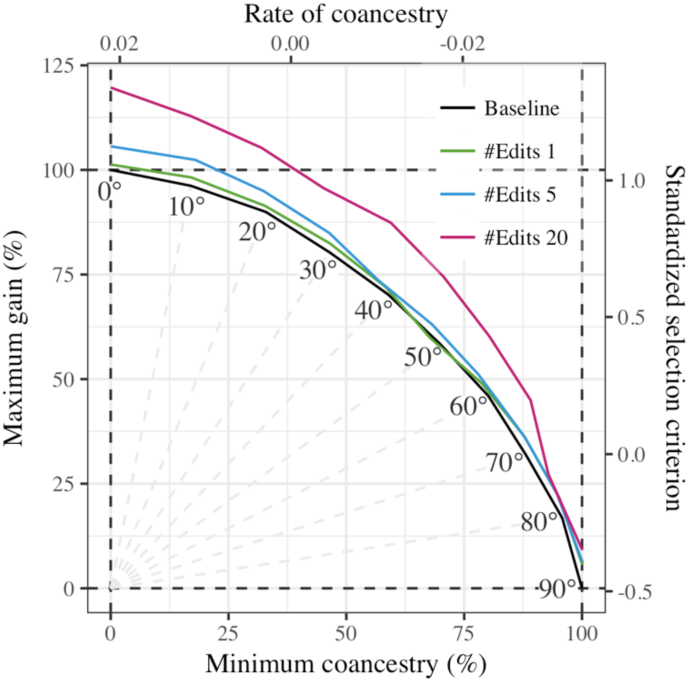
Trade-off between genetic gain and group coancestry and its modification with genome editing

## 5 CONCLUSION

In this paper, we have described the AlphaMate program that optimises selection, maintenance of diversity, and mate allocation in breeding programs. The program enables both animal and plant breeding programs to be more optimal and facilitates new research opportunities.

Conflict of Interest: none declared.

